# Robustness of NeuroMark-Derived Functional Networks to fMRI Spatial Normalization Across the Human Lifespan

**DOI:** 10.64898/2026.04.24.720677

**Authors:** Zening Fu, Sarah Shultz, Armin Iraji, Vince D. Calhoun

## Abstract

NeuroMark is a fully automated hybrid independent component analysis (ICA) framework designed to extract functional network features that are individually resolved and comparable across different cohorts. By integrating a reliable spatial template with spatially constrained ICA that adapts to each scan, NeuroMark retains the advantages of data-driven decomposition while avoiding limitations of fixed region-of-interest approaches. NeuroMark typically employs direct spatial normalization of fMRI data to a standardized adult EPI template; it remains unclear whether this approach is optimal for populations whose anatomy differs substantially from that of adults. We evaluated two normalization strategies in three large datasets spanning infancy, development, and aging: (1) direct normalization to the adult EPI template (EPInorm), and (2) normalization using an age-specific anatomical T1 template followed by transformation to the adult EPI template (T1toEPInorm). Across all cohorts, average intrinsic connectivity networks derived from EPInorm and T1toEPInorm exhibited very high spatial correspondence (mean ± SD: 0.9966 ± 0.0012 in infants; 0.9947 ± 0.0019 in development; 0.9963 ± 0.0012 in aging). The individual level also showed high similarity, though time courses showed slightly higher consistency than spatial maps (average correlations for time courses: 0.7990–0.9931; average correlations for spatial maps: 0.6879–0.9131). Functional network connectivity (FNC) measures were extremely well preserved across scans (95% of FNC with r > 0.9374 in infants; r > 0.8670 in developmental cohorts; r > 0.9219 in aging), demonstrating the robustness of NeuroMark features to different normalization strategies. Together, these results indicate that NeuroMark yields highly stable functional network features irrespective of whether an age-specific intermediate registration step is incorporated. NeuroMark, along with direct normalization to the adult EPI template, thus provides a robust, efficient, and harmonizable approach for large-scale, multisite, and lifespan neuroimaging studies, facilitating broad comparability across datasets while avoiding potential biases introduced by using multiple age-specific templates within a single study.

## Introduction

NeuroMark is an advanced independent component analysis (ICA)-based framework (Du et al., 2020) designed to delineate intrinsic connectivity networks (ICNs) and extract network-specific functional neuroimaging features, thereby contributing to a deeper understanding of large-scale functional brain organization. Methodologically, NeuroMark occupies a unique position at the intersection of atlas-based (Faria et al., 2012; Rolls et al., 2020) and data-driven (Beckmann et al., 2009; Correa et al., 2007; Smith et al., 2014) analytic strategies, two predominant approaches for parcellating the brain (Calhoun, 2025). By integrating a high-quality spatial reference template with spatially constrained ICA performed at the individual-subject level, NeuroMark enables the derivation of functional features that are directly comparable across subjects, studies, and datasets. This enhances methodological reproducibility, enables automated biomarker discovery pipelines, and confers many of the strengths associated with atlas-based frameworks (Fu et al., 2024). At the same time, its data-driven decomposition allows the spatial parcellation to adapt to each scan, overlap, and yield continuous values at each voxel rather than a fixed average, thereby overcoming key limitations of fixed ROIs, which may not accurately reflect the underlying functional architecture. This scan-specific adaptation enables a more precise characterization of inter-individual variability, ensuring that individual-level feature extraction remains completely independent of group-level modeling and thereby mitigates concerns about training-testing leakage in large-scale studies (Du and Fan, 2013). Contemporary neuroscience is increasingly defined by the challenges of generalizability and reproducibility in population-level research (Marek et al., 2022), with large-scale neuroimaging consortia now providing unprecedented sample sizes for cross-dataset and cross-cohort investigations (Bycroft et al., 2018; Luciana et al., 2018; Wu-Minn, 2017). Within this context, NeuroMark is well positioned to support robust multisite analyses, as its only principal prerequisite is spatial normalization of functional magnetic resonance imaging (fMRI) data to the standard adult Montreal Neurological Institute (MNI) space (Tzourio-Mazoyer et al., 2002). This requirement enables consistent representation of large-scale functional networks across heterogeneous demographic groups and imaging datasets, facilitating high-throughput, harmonized analyses aligned with current large-scale neuroimaging research priorities.

Spatial normalization of fMRI data is a fundamental component of neuroimaging preprocessing. It is implemented across all major software toolkits, with numerous established strategies available for warping echo planar imaging (EPI) data into standardized anatomical spaces (Collins et al., 1994; Mazziotta et al., 2001). Within most NeuroMark applications, a commonly used approach—EPInorm—applies an affine transformation followed by nonlinear registration of each subject’s EPI image directly to an adult EPI template in Montreal Neurological Institute (MNI) space (Friston et al., 1995, 1999; Klein et al., 2009). This method explicitly models the nonlinear spatial distortions inherent in EPI acquisitions; however, it may also introduce the risk of overcorrection, particularly in regions where distortions are severe or spatial correspondences are ambiguous. An alternative strategy, referred to as T1toEPInorm, incorporates high-resolution structural information to guide the normalization process. In this approach, a subject-specific or population-derived T1-weighted structural image is used to estimate the affine and nonlinear warping fields necessary for mapping data to MNI space (Friston et al., 1995; Tzourio-Mazoyer et al., 2002). Leveraging a higher spatial resolution anatomical image can improve anatomical fidelity and increase registration accuracy, because the transformation is informed by tissue boundaries and subject- or cohort-specific morphology. This strategy has been widely adopted in large population-level initiatives, including the UK Biobank (Littlejohns et al., 2020) and the Human Connectome Project (Glasser et al., 2013), and has demonstrated reliable performance for spatial normalization of EPI data. Despite the trade-off that structural-based normalization cannot correct for geometric distortions in the EPI data, there is increasing recognition of the need for age- or population-specific high-resolution brain templates (Yushkevich et al., 2009; Zhao et al., 2016), as templates derived from young healthy adults do not fully represent neuroanatomical variation across development, adulthood, and aging (Machilsen et al., 2007). Maturational changes in gray matter morphology, cortical thickness, and white matter microstructure, and brain shape (Ge et al., 2002; Lemaître et al., 2005; Lenroot and Giedd, 2006) across the lifespan suggest that normalization based solely on adult reference templates may introduce bias or misalignment in studies of infants, children, or older adults (Fonov et al., 2011; Hoeksma et al., 2005; Ridwan et al., 2021; Wilke et al., 2002a). Such work has emphasized the importance of population-matched templates to improve alignment accuracy and reduce downstream analytical confounds.

There are some limitations, though; for example, if a single study uses multiple age-specific templates, this can inflate age-related changes. Although some other evidence suggests that EPInorm can support voxel-level analyses that perform comparably or even outperform approaches incorporating structural images, even followed by distortion correction (Calhoun et al., 2017), it remains unknown whether this standard direct-registration strategy is fully robust when applied to diverse demographic cohorts within the NeuroMark framework. Moreover, it remains unresolved whether incorporating an intermediate, age-specific high-resolution structural normalization step would materially influence the intrinsic connectivity networks or downstream network-level biomarkers extracted by NeuroMark. A systematic evaluation of these questions is therefore required to determine whether the normalization strategy affects NeuroMark network characterization across differing age populations.

The objective of this study is to comprehensively evaluate the performance of two common normalization strategies—the EPInorm approach and the T1toEPInorm approach in NeuroMark analyses across different age groups. To accomplish this, we leveraged three large-scale neuroimaging datasets that collectively capture critical stages of the human lifespan: infancy (birth to approximately nine months), childhood through young adulthood (five to 21 years), and middle to late adulthood (36 to 100 years). These complementary cohorts allow assessment of normalization performance across periods characterized by substantial neuroanatomical and functional reorganization. The primary evaluation metric was the correlation coefficient between NeuroMark features derived using the two normalization schemes, providing a quantitative measure of the degree of concordance between EPInorm and T1toEPInorm. While this work focuses on the NeuroMark Functional 1.0 template (the original and currently most widely used template in NeuroMark studies) (Du et al., 2020), the findings are expected to generalize across other NeuroMark functional templates (Fouladivanda et al., 2024; Fu et al., 2024). To assess the robustness and consistency of normalization choices, we examined three popularly used NeuroMark-derived features: (1) spatial maps of intrinsic connectivity networks, (2) network time courses, and (3) functional network connectivity (FNC). Comparisons were performed at both the group and individual levels to fully characterize the impact of normalization choice on derived neurofunctional representations. To our knowledge, this represents the first systematic investigation into whether population-tailored anatomical registration— designed to account for age-related differences in brain morphology—is necessary when applying NeuroMark to cohorts whose neuroanatomy deviates substantially from that of young adults. The findings provide insight into the stability and generalizability of NeuroMark-derived features across the lifespan and inform best practices for preprocessing in large-scale lifespan neuroimaging studies.

## Methods

### Analysis Flowchart

**Fig. 1** presents the analytical workflow to evaluate whether the choice of spatial normalization strategy during fMRI preprocessing influences downstream outputs within the NeuroMark framework. After initial preprocessing using a standardized pipeline, fMRI data were spatially normalized to MNI space using either the EPInorm or T1toEPInorm procedure. NeuroMark was subsequently applied to each normalized dataset to estimate individualized ICNs and their corresponding time courses, from which FNC matrices were derived. These analyses were conducted across three large, developmentally distinct cohorts—infants (0–9 months), children and adolescents (5–21 years), and aging adults (36–100 years)—thereby encompassing major phases of neurodevelopmental maturation and age-related neurodegeneration. Three principal NeuroMark-derived features were examined: spatial ICN maps, temporal dynamics of ICN time courses, and inter-network FNC. Comparisons between normalization strategies were performed to assess the consistency of these features and to determine whether the choice of spatial normalization systematically alters NeuroMark’s characterization of large-scale functional brain organization.

**Figure 1.**
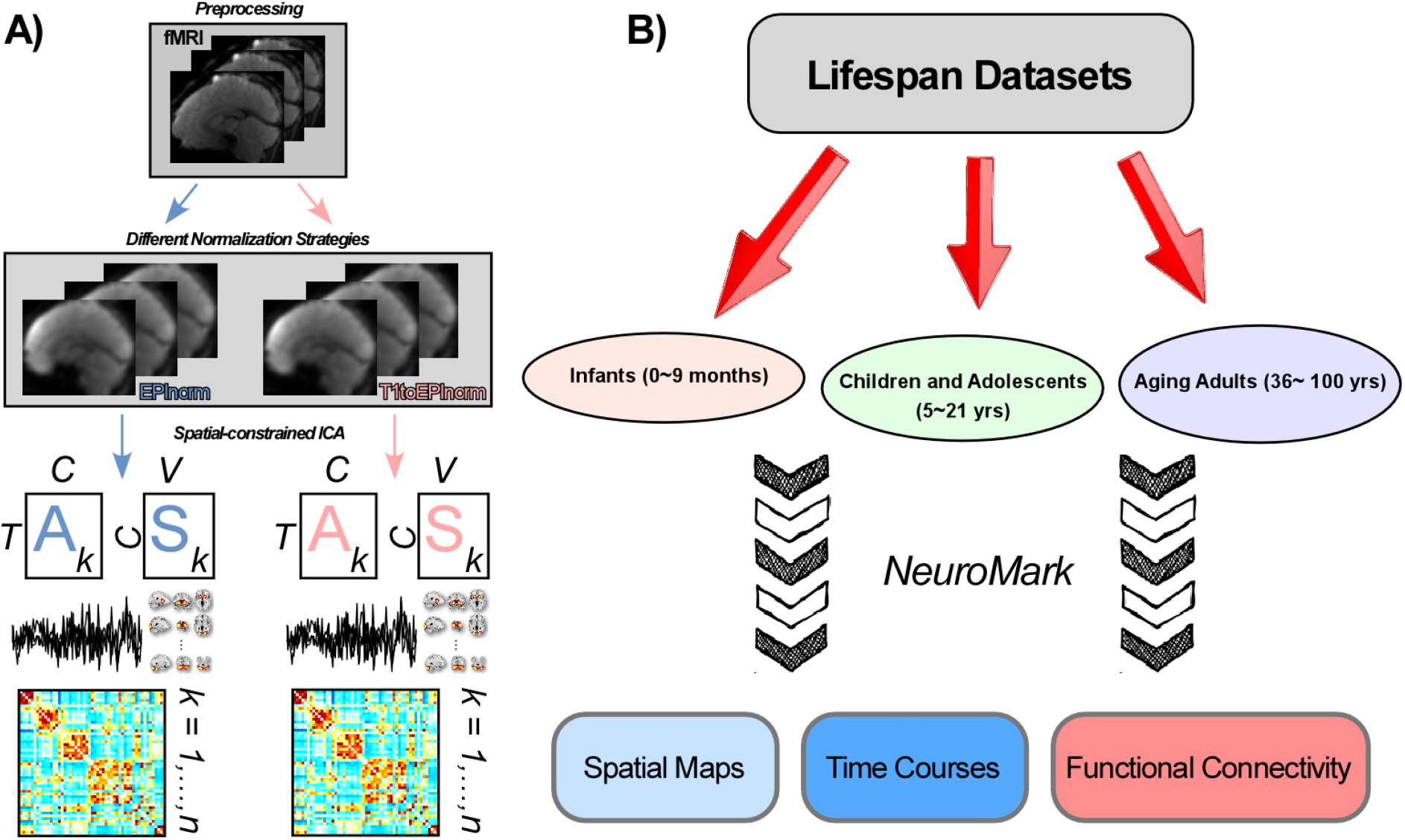
Evaluation of spatial normalization strategies for NeuroMark feature extraction across lifespan neuroimaging cohorts. **(A)** Preprocessed fMRI data are spatially normalized to MNI space using either EPInorm or T1toEPInorm, after which individual-level features are reconstructed with reference to the NeuroMark functional 1.0 template. Template information–guided back-reconstruction procedures are applied to each scan to obtain feature estimates that are directly comparable across normalization strategies. **(B)** These analyses are conducted on three large-scale datasets representing distinct developmental stages: infancy (0–9 months), childhood through young adulthood (5–21 years), and middle to late adulthood (36–100 years). Three commonly examined NeuroMark-derived feature domains are assessed: intrinsic connectivity network spatial maps, network time courses, and functional (network) connectivity.

### Datasets Spanning Distinct Age Cohorts

The present study sought to determine whether directly normalizing fMRI data from individuals across markedly different developmental stages to a standard adult template introduces systematic biases in the estimation of intrinsic neural network architecture. To address this question, we analyzed three large-scale resting-state fMRI datasets spanning major periods of human brain maturation and aging: infancy (birth to nine months), the developmental period (five to 21 years), and aging adulthood (36 to over 100 years). Together, these cohorts enabled a comprehensive evaluation of spatial normalization effects across the lifespan, encompassing phases of rapid neurodevelopment, relative structural stability, and age-related neuroanatomical decline.

The infant cohort comprised data from the Neuroimaging of Infants at High- and Low-Risk for ASD (NI-HLA) and the ACE Center 2017: Project 3 – Neuroimaging (ACE) studies (Feinberg et al., 2010; Moeller et al., 2010), both conducted at the Center for Systems Imaging at the Emory University School of Medicine. Infants were scanned longitudinally during natural sleep at multiple time points between birth and nine months, allowing characterization of early-emerging functional network structure during a period of rapid cortical and subcortical development. The developmental cohort was drawn from the Human Connectome Project-Development (HCP-D) (Somerville et al., 2018), a large publicly available dataset containing resting-state fMRI from 652 typically developing children, adolescents, and young adults aged 5–21 years. This dataset captures the transition from childhood through adolescence into early adulthood, encompassing substantial changes in cortical thickness, white matter maturation, and functional network organization. The aging cohort was obtained from the Human Connectome Project-Aging (HCP-A) (Bookheimer et al., 2019), including resting-state fMRI from 725 neurologically healthy adults aged 36 to beyond 100 years. This dataset provides a broad sampling of middle age through late life, permitting examination of normalization effects during periods associated with brain atrophy, reduced network segregation, and other age-related neurobiological alterations. Further details regarding these datasets are presented in the Supplementary Materials.

### Imaging Preprocessing

We preprocessed fMRI datasets using established pipelines implemented with the FMRIB Software Library (FSL v6.0) and Statistical Parametric Mapping (SPM12), executed within the MATLAB 2024b environment. Preprocessing followed standard best practices for resting-state fMRI and included several steps designed to improve signal stability and imaging alignment. First, the initial ten volumes of each scan were discarded to allow for signal equilibrium and reduce transient magnetization effects. Rigid-body realignment was then applied to correct for head motion, followed by distortion correction to compensate for field inhomogeneities and geometric warping commonly encountered in EPI acquisitions. Slice-timing correction was performed to account for temporal differences in slice acquisition. Subsequently, each dataset was spatially normalized to the MNI standard space using one of two normalization strategies: EPInorm or T1toEPInorm. Normalized images were resampled to an isotropic resolution of 3 × 3 × 3 mm^3^ and spatially smoothed with a Gaussian kernel of 6 mm full width at half maximum (FWHM) to enhance signal-to-noise characteristics and spatial correspondence across scans. Additional methodological details and parameter specifications are provided in the Supplementary Materials.

### Normalization to MNI Space

This study assessed two commonly used spatial normalization methods within the preprocessing framework: EPInorm and T1toEPInorm. EPInorm is a single-step procedure where EPI images are directly registered to the standard young-adult EPI template provided with SPM (Penny et al., 2011). This method has been widely employed in our previous ICA-based studies. T1toEPInorm uses a two-stage normalization process. In this approach, functional data are first aligned with an age-appropriate structural T1-weighted template and then transformed into the adult MNI EPI space. This strategy takes advantage of age-specific anatomical information to enhance warping accuracy, reflecting a commonly adopted practice in large-scale neuroimaging pipelines, such as those utilized by the Human Connectome Project and other population-level imaging repositories.

For the infant cohort, we employed densely sampled age-specific T1 templates from the UNC/UMN Baby Connectome Project (BCP) (Chen et al., 2022), which capture dynamic brain development during infancy. For the developmental cohort, we utilized unbiased pediatric T1 atlases (ages 4.5–18.5 years) from the NeuroImaging & Surgical Technologies Lab at McGill University (Fonov et al., 2011, 2009), constructed to represent normative brain morphology across this age range. For the aging cohort, we adopted the Multichannel Illinois Institute of Technology and Rush University Aging (MIITRA) atlas (Wu et al., 2023), which comprises a high-quality T1 template generated through advanced multimodal template-construction techniques using data from 400 cognitively healthy older adults. Two versions of preprocessed fMRI data were thus produced for each cohort, corresponding to the EPInorm and T1toEPInorm normalization, and subsequently analyzed within the NeuroMark framework for comparative evaluation.

### NeuroMark: Functional Network Feature Extraction

We applied the NeuroMark framework with the functional 1.0 template to the preprocessed fMRI data within the Group ICA of fMRI Toolbox (GIFT; http://trendscenter.org/software/gift). NeuroMark represents an advancement over both traditional atlas-based parcellation and unconstrained ICA by incorporating spatially constrained decomposition driven by large-sample, population-derived functional priors. This strategy enables consistent and biologically meaningful extraction of subject-level ICNs and their associated time courses, mapped to 53 predefined functional network priors distributed across seven canonical domains: subcortical (SC), auditory (AUD), sensorimotor (SM), visual (VS), cognitive control (CC), default mode (DM), and cerebellar (CB).

Three primary NeuroMark-derived features were evaluated: **(1)** ICN spatial maps, representing voxelwise network expression; **(2)** network-specific time courses, indexing the temporal dynamics of ICN activity; and **(3)** FNC, a higher-order measure characterizing inter-network coupling analogous to conventional resting-state functional connectivity. Before FNC estimation, ICN time courses underwent an additional series of post-processing steps to minimize residual non-neuronal variance. Specifically, we applied **(i)** polynomial detrending (linear, quadratic, and cubic) to remove low-frequency drifts, **(ii)** regression of six rigid-body motion parameters and their first derivatives, **(iii)** automated detection and exclusion of extreme outlier time points, and **(iv)** temporal band-pass filtering (0.01–0.15 Hz) to preserve physiologically relevant low-frequency fluctuations. FNC matrices were then computed as pairwise Pearson correlation coefficients among the denoised ICN time courses.

### Similarity Correlation Analysis between Normalization Strategies

To systematically assess the degree of concordance between imaging features derived from different spatial normalization strategies, we quantified feature similarity using correlation-based metrics. For each dataset, we first computed correlations between group-level spatial maps for each ICN. Group-level maps were obtained by averaging network-specific spatial maps across all scans within the dataset, providing a population-level representation of network organization. To further evaluate consistency at the individual level, spatial map correlations were also computed for each scan by comparing the ICN maps produced under the two normalization approaches. In parallel, we examined whether the network fluctuations were preserved across normalization strategies by calculating the correlation between network time courses for each scan. This analysis aimed to determine whether NeuroMark consistently captures intrinsic neural fluctuations across alignment pathways. We further assessed FNC similarity analogously, with correlations computed at both the group and individual-scan levels. As FNC is a central measure in brain-wide association studies (Marek et al., 2022), we additionally calculated the correlation between pairwise FNC across scans to evaluate the robustness of cross-scan/subject variability of connectivity features under differing normalization conditions.

## Results

### Infant Cohorts show High Consistency of NeuroMark-Derived Spatial Networks and Temporal Fluctuations between Normalization Strategies

The infant cohort included 500 resting-state fMRI scans from 170 participants, as shown in **Table I**. We first derived the group-level spatial maps of ICNs across scans and assessed their correspondence under the two normalization strategies. As shown in **Fig. 2A**, the spatial maps obtained from both approaches exhibited high similarity across all seven functional domains in the infant cohort. The correlation coefficients between averaged spatial maps of corresponding networks are summarized in **Fig. 2B**, demonstrating extremely high similarity (all correlations > 0.99; mean ± SD = 0.9966 ± 0.0012, across ICNs).

**Table I.** Basic Demographics of Lifespan Datasets.

| Basic Demographics | Infant Cohorts | Developmental Cohorts | Aging Cohorts |
| --- | --- | --- | --- |
| Total Subject | 170 | 652 | 725 |
| Scans | 500 | 2,513 | 2,890 |
| Age (month) [scans] | $3.36 \pm 1.83$ | $175.80 \pm 46.79$ | $724.07 \pm 188.29$ |
| Gender (F/M) [scans] | 222/278 | 1351/1162 | 1619/1271 |
| Mean FD [scans] | $0.28 \pm 0.30$ | $0.16 \pm 0.08$ | $0.20 \pm 0.08$ |

**Figure 2.**
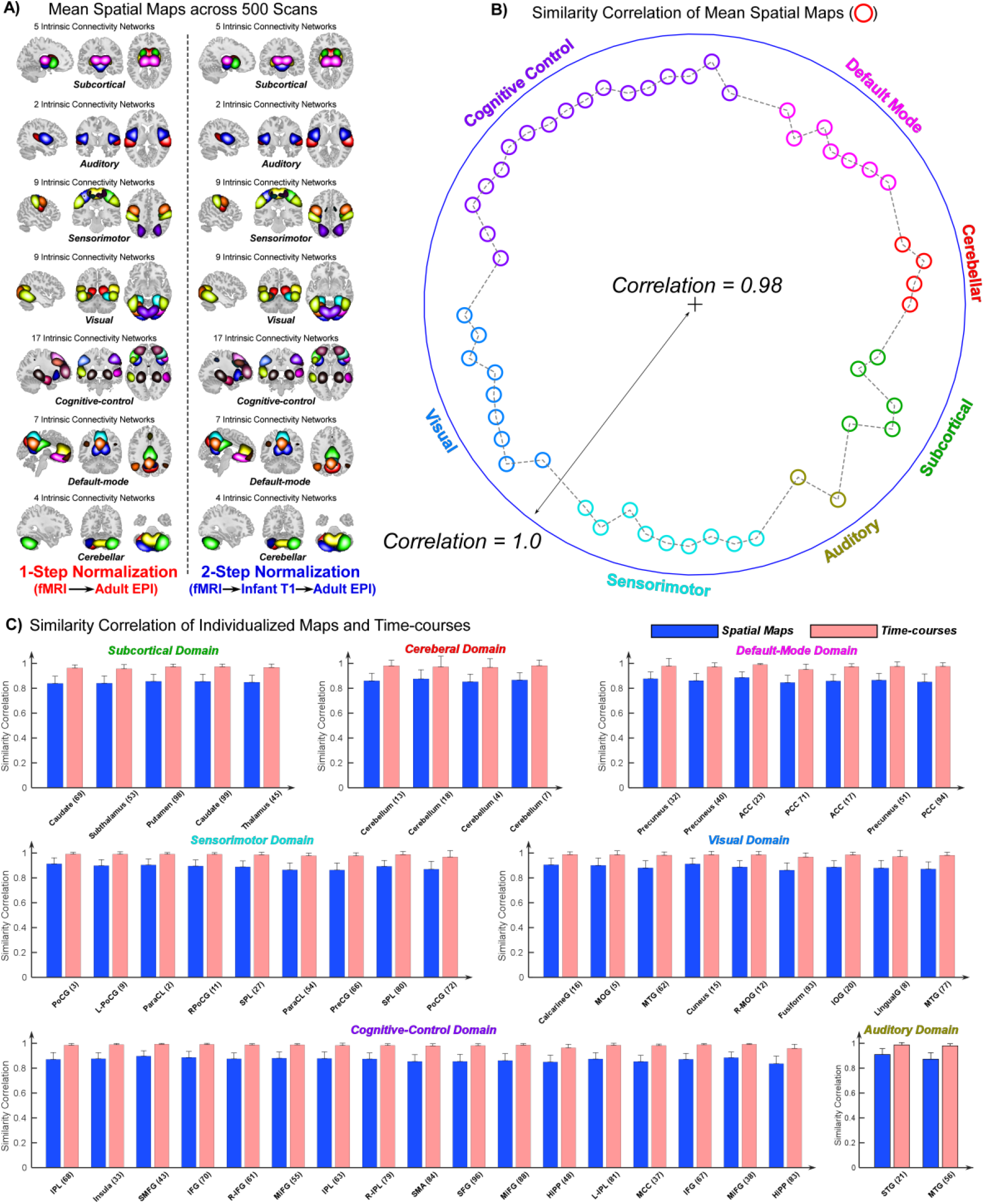
Comparison of spatial maps and time courses of infant networks obtained under different normalization strategies. **(A)** Spatial maps were averaged across 500 scans and aggregated within each of the seven functional domains. High correspondence was observed between networks normalized directly to the adult EPI template and those normalized first to an infant-specific template and subsequently to the adult template, indicating that direct normalization to the adult template does not substantially alter network spatial configurations. **(B)** Pearson correlation coefficients between mean spatial maps of corresponding networks across strategies. **(C)** Pearson correlation coefficients between individualized spatial maps and time courses across strategies. Bar plots represent mean correlations averaged across 500 scans, with error bars denoting standard deviations. Although time courses exhibited greater inter-strategy similarity than spatial maps, all mean correlation values exceeded 0.83, demonstrating strong consistency of network features across normalization approaches.

We further computed correlations between spatial maps and time courses of corresponding networks for each scan, yielding a network-specific spatial correlation and a network-specific temporal correlation. The results, summarized in **Fig. 2C**, show the mean and standard deviations of correlations across 500 scans, with spatial similarity represented by blue bars and temporal similarity represented by pink bars. Among functional domains, the auditory domain exhibited the highest spatial similarity (mean correlation = 0.8912), whereas the subcortical domain showed the lowest (mean correlation = 0.8479). For time courses, the sensorimotor domain demonstrated the highest similarity (mean correlation = 0.9843), while the subcortical domain again showed the lowest (mean correlation = 0.9670). At the network level, the postcentral gyrus (PoCG, 3) demonstrated the highest spatial consistency (mean ± SD = 0.9131 ± 0.0483, across scans), whereas the hippocampus (HiPP, 83) exhibited the lowest spatial similarity (mean ± SD = 0.8361 ± 0.0607, across scans). On the other hand, the superior medial frontal gyrus (SMFG, 43) showed the greatest temporal consistency (mean ± SD = 0.9931 ± 0.0066, across scans), while the posterior cingulate cortex (PCC, 71) showed the lowest (mean ± SD = 0.9510 ± 0.0436, across scans). Overall, time courses (mean correlation = 0.9800) demonstrated higher similarity than spatial maps (mean correlation = 0.8730). Nevertheless, all networks yielded mean correlations exceeding 0.83 for both features, indicating high consistency between normalization strategies.

### Developmental Cohorts show High Consistency of NeuroMark-Derived Spatial Networks and Temporal Fluctuations between Normalization Strategies

The developmental cohort comprised 2,513 resting-state fMRI scans from 652 children and adolescents. Findings for the developmental cohort are presented in **Fig. 3**. Consistent with the infant cohort, spatial maps derived using the two normalization strategies exhibited high spatial similarity across all seven functional domains (**Fig. 3A**). Correlation analysis of the averaged spatial maps confirmed this correspondence, with all coefficients exceeding 0.98 (mean ± SD = 0.9947 ± 0.0019, across ICNs; **Fig. 3B**).

**Figure 3.**
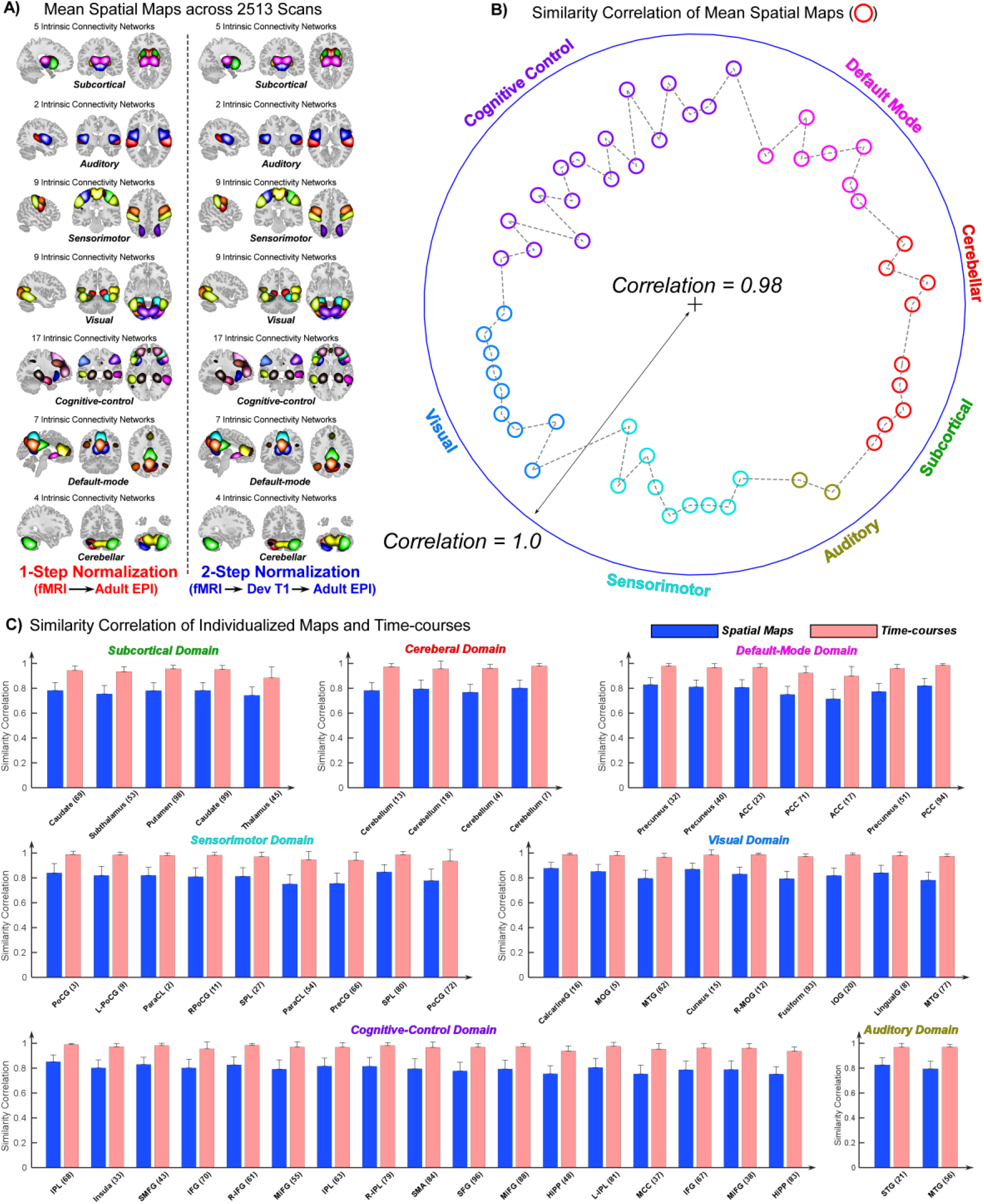
Comparison of spatial maps and time courses of developmental networks obtained under different normalization strategies. **(A)** Spatial maps were averaged across 2,513 scans and aggregated within each of the seven functional domains. High correspondence was observed between networks normalized directly to the adult EPI template and those normalized first to a child/adolescent template and subsequently to the adult template, indicating that direct normalization to the adult template does not substantially alter network spatial configurations. **(B)** Pearson correlation coefficients between mean spatial maps of corresponding networks across strategies. **(C)** Pearson correlation coefficients between individualized spatial maps and time courses across strategies. Bar plots represent mean correlations averaged across 2,513 scans, with error bars denoting standard deviations. Although time courses exhibited greater inter-strategy similarity than spatial maps, all mean correlation values exceeded 0.71, demonstrating strong consistency of network features across normalization approaches.

For the scan-level analysis, **Fig. 3C** shows the mean and standard deviations of correlations for each network across 2,513 scans. Among functional domains, the visual domain exhibited the highest spatial similarity (mean correlation = 0.8297), whereas the subcortical domain showed the lowest (mean correlation = 0.7686). Similarly, time courses were most consistent in the visual domain (mean correlation = 0.9810) and least consistent in the subcortical domain (mean correlation = 0.9339). At the network level, the calcarine gyrus (CalcarineG, 16) demonstrated the strongest spatial correspondence (mean ± SD = 0.8781 ± 0.0471, across scans), while the anterior cingulate cortex (ACC, 17) exhibited the weakest (mean ± SD = 0.7163 ± 0.0771, across scans). For time courses, the inferior parietal lobule (IPL, 68) showed the greatest similarity (mean ± SD = 0.9907 ± 0.0080, across scans), whereas the thalamus (45) showed the lowest (mean ± SD = 0.8837 ± 0.0883, across scans). Again, time courses (mean correlation = 0.9654) demonstrated greater cross-strategy similarity than spatial maps (mean correlation = = 0.7995). Although the developmental cohort exhibited slightly lower correspondence compared to the infant data, all networks maintained mean correlations above 0.71 for both features, underscoring the robustness of network estimation across normalization strategies.

### Aging Cohorts show High Consistency of NeuroMark-Derived Spatial Networks and Temporal Fluctuations between Normalization Strategies

The aging cohort comprised 2,890 resting-state fMRI scans from 725 older adults. The results from the aging cohort were consistent with those observed in the infant and developmental populations. As shown in **Fig. 4A**, spatial maps derived using the two normalization strategies exhibited high similarity across all seven functional domains. Correlation analysis of the averaged spatial maps confirmed extremely high correspondence between normalization strategies, with all coefficients exceeding 0.99 (mean ± SD = 0.9963 ± 0.0012, across ICNs; **Fig. 4B**).

**Figure 4.**
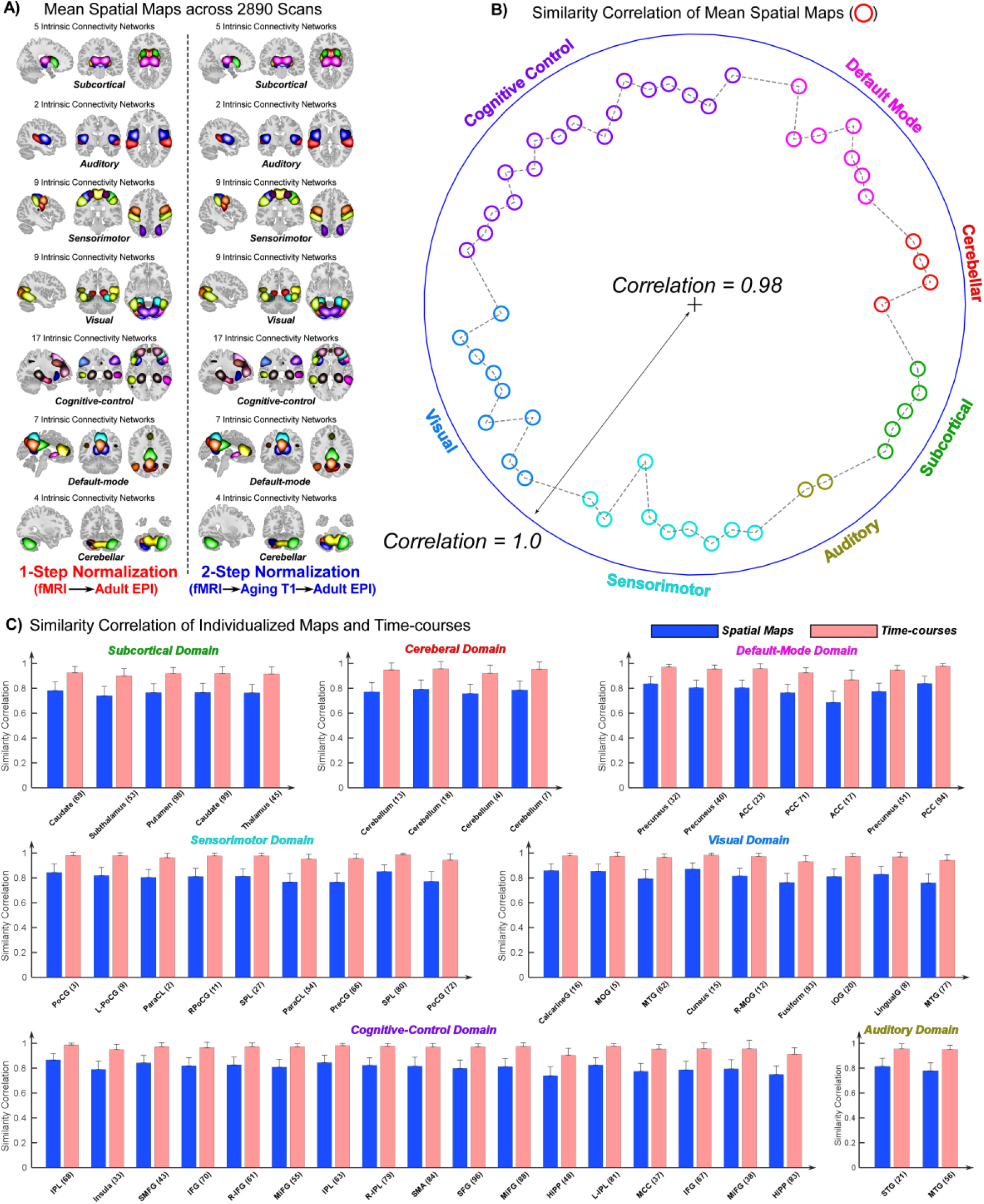
Comparison of spatial maps and time courses of aging networks obtained under different normalization strategies. **(A)** Spatial maps were averaged across 2,890 scans and aggregated within each of the seven functional domains. High correspondence was observed between networks normalized directly to the adult EPI template and those normalized first to an aging adult template and subsequently to the adult template, indicating that direct normalization to the adult template does not substantially alter network spatial configurations. **(B)** Pearson correlation coefficients between mean spatial maps of corresponding networks across strategies. **(C)** Pearson correlation coefficients between individualized spatial maps and time courses across strategies. Bar plots represent mean correlations averaged across 2,890 scans, with error bars denoting standard deviations. Although time courses exhibited greater inter-strategy similarity than spatial maps, all mean correlation values exceeded 0.68, demonstrating strong consistency of network features across normalization approaches.

Scan-level analyses further supported these findings. As illustrated in **Fig. 4C**, the bar plots show mean and standard deviations of correlations across 2,890 scans. At the domain level, the visual domain exhibited the highest spatial similarity (mean correlation = 0.8181), whereas the subcortical domain showed the lowest (mean correlation = 0.7623). For time courses, the sensorimotor domain demonstrated the greatest similarity (mean correlation = 0.9672), while the subcortical domain again showed the lowest (mean correlation = 0.9154). Network-level results revealed that the cuneus (15) exhibited the strongest spatial correspondence (mean ± SD = 0.8720 ± 0.0501, across scans), whereas the ACC (17) showed the weakest (mean ± SD = 0.6879 ± 0.0894, across scans). For time courses, the IPL (68) demonstrated the highest similarity (mean ± SD = 0.9871 ± 0.0142, across scans), while the ACC (17) again showed the lowest (mean ± SD = 0.8698 ± 0.0788, across scans). Overall, time courses (mean correlation = 0.9555) demonstrated higher similarity than spatial maps (mean correlation = 0.7990) across functional domains. Importantly, all networks yielded correlation values above 0.68 for both features, indicating that direct normalization to the adult EPI template does not introduce systematic bias into NeuroMark-derived features in the aging cohort.

### NeuroMark-Derived FNC Robust to Normalization Strategies across Lifespan

We assessed the similarities of FNC, a key feature investigated in ICA-based studies. **Fig. 5A** (upper panel) displays the group-level FNC matrix averaged across 500 infant scans for each normalization strategy. No structural differences were observed between the connectivity profiles, suggesting that direct normalization of infant data to the adult EPI template does not induce significant systematic distortions in FNC. The scatter plot in **Fig. 5A** further demonstrates a near-perfect correlation between the group-level FNC matrices from the two normalization approaches (r = 0.9974, p < 10^−10^). We next quantified similarity at the individual scan level. The plot in **Fig. 5A** shows correlations between individualized FNC matrices between normalization strategies, where each asterisk represents a single scan. Infant data exhibited exceptionally high similarity (mean ± SD = 0.9845 ± 0.0179, across scans), indicating strong preservation of FNC at the scan level. Comparable findings were observed in developmental and aging cohorts (**Figs. 5B** and **5C**), with mean similarity values of 0.9649 ± 0.0261 and 0.9785 ± 0.0196, across scans, respectively.

**Figure 5.**
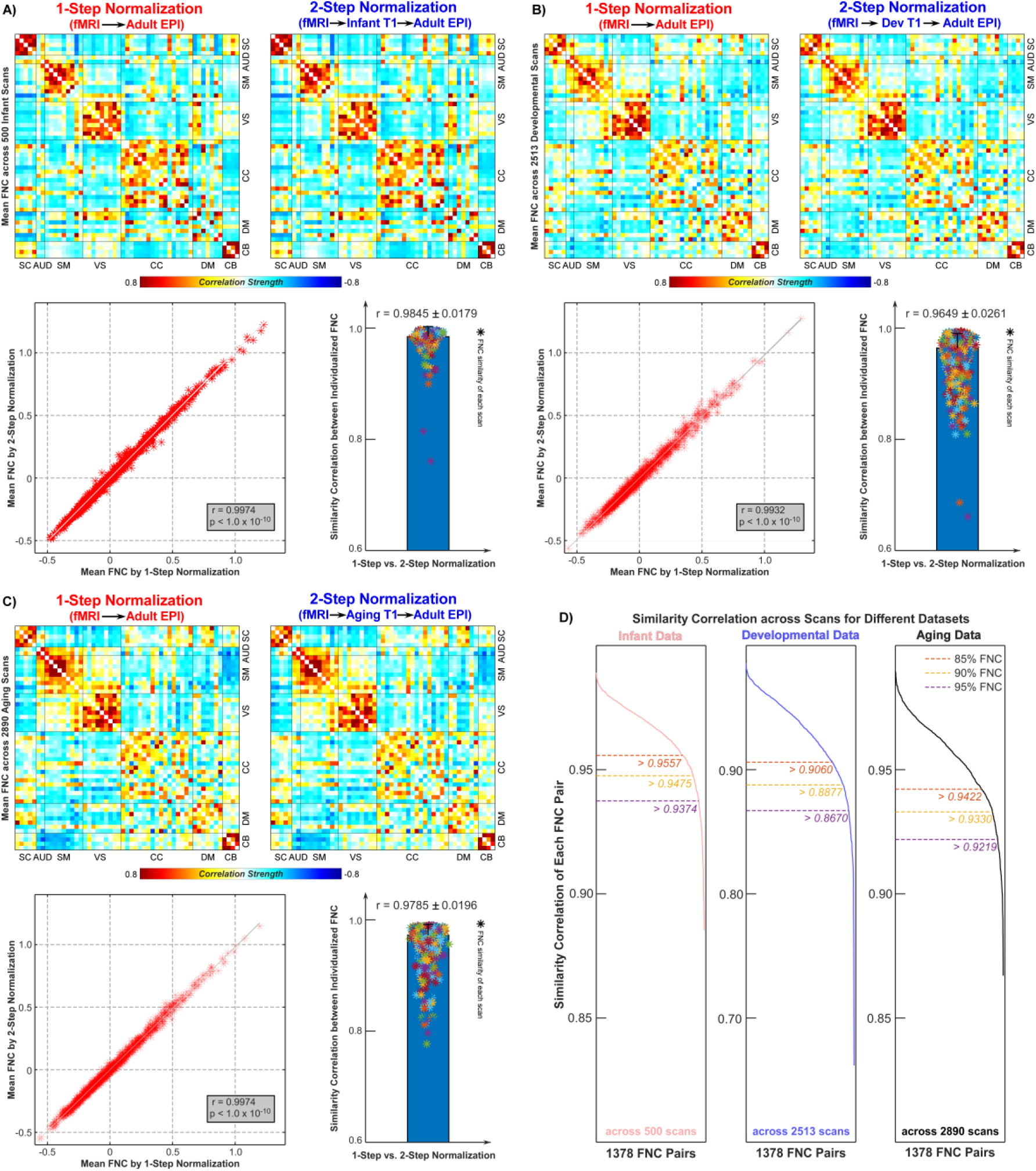
Comparison of functional network connectivity (FNC) derived from datasets processed using different normalization strategies. **(A)** FNC results for the infant cohort. Connectivity matrices were averaged across 500 scans and organized according to seven functional domains. High correspondence is observed between FNC profiles obtained under the two normalization strategies, indicating that direct normalization to the adult EPI template does not substantially alter network connectivity patterns. The scatter plot depicts the correlation between mean FNC matrices from the two strategies, demonstrating an exceptionally high similarity (r = 0.9974, p < 10^−10^). The bar plot illustrates correlations between individualized FNC matrices, confirming similarly high consistency at the single-scan level (r = 0.9845 ± 0.0179). **(B)** FNC results for the developmental cohort. **(C)** FNC results for the aging cohort. **(D)** Correlations for each FNC pair across scans between the two normalization strategies. Percentile thresholds (85%, 90%, and 95%) are highlighted, revealing that the majority of FNC pairs exhibit highly consistent inter-scan variation (r > 0.9).

We further evaluated cross-scan variability for each FNC pair, which is a crucial factor in both univariate and multivariate brain-wide association analyses in neuroimaging studies. Specifically, for each FNC pair, we had an N × 1 vector of estimates from EPInorm and an N × 1 vector of estimates from T1toEPInorm (N is the number of scans), and then computed the correlation between these two vectors. In total, 1378 FNC pairs were analyzed, reflecting whole-brain connectivity among 53 NeuroMark networks. Results are summarized in **Fig. 5D**. For the infant cohort, 85%, 90%, and 95% of FNC pairs exhibited correlations exceeding 0.9557, 0.9475, and 0.9374, respectively, with a minimum correlation of r = 0.8853 (p < 10^−10^). For the developmental cohort, the corresponding thresholds were 0.9060, 0.8877, and 0.8670, with a minimum correlation of r = 0.6671 (p < 10^−10^). For the aging cohort, thresholds were 0.9422, 0.9330, and 0.9219, with a minimum correlation of r = 0.8670 (p < 10^−10^). Our findings suggest that the results of FNC-behavior association studies may remain consistent regardless of the normalization strategies employed.

## Discussion

These convergent findings demonstrate that NeuroMark exhibits strong robustness and cross-cohort generalizability across populations spanning distinct stages of the lifespan, independent of the age-specific normalization strategy employed. Core NeuroMark-derived features—including spatial ICN profiles, intrinsic temporal dynamics, and inter-network functional connectivity—remain highly stable across normalization approaches. This consistency persists whether fMRI data are registered directly to the adult MNI template or normalized via age- or population-specific structural references, indicating that NeuroMark reliably recovers intrinsic functional architecture even in cohorts whose neuroanatomy diverges substantially from that of young adults.

ICA is a powerful, data-driven framework for identifying intrinsic functional networks and other neuroimaging features across modalities (Calhoun et al., 2009). However, conventional ICA implementations can produce solutions that vary across runs and datasets, complicating cross-study reproducibility and limiting direct comparisons in large-scale population neuroscience. As neuroimaging increasingly emphasizes multi-cohort validation and cross-disorder characterization, reliable and standardized feature extraction has become essential (Miłkowski et al., 2018). Many neuropsychiatric and neurodegenerative disorders exhibit both overlapping and disorder-specific neural alterations, underscoring the value of consistent analytic frameworks for differentiating shared and unique network abnormalities (Du et al., 2021; Fu et al., 2019, 2020, 2021b). The emergence of large public neuroimaging consortia has further highlighted the need for ICA approaches that yield stable and comparable components across studies and clinical populations. Therefore, we developed the NeuroMark framework, which integrates data-driven decomposition with standardized templates. By anchoring component estimation to predefined functional priors, NeuroMark ensures one-to-one correspondence of imaging features across individuals, studies, and datasets, while the ICA-based optimization retains subject-specific variability critical for characterizing individual differences. This hybrid strategy combines the strengths of atlas-driven and unconstrained ICA approaches, enabling reliable extraction of spatial networks, temporal dynamics, and functional network connectivity measures. NeuroMark has demonstrated strong utility in both normative populations and diverse clinical cohorts, providing a robust platform for identifying reproducible functional biomarkers and supporting large-scale comparisons across brain disorders (Dhamala et al., 2023; Fu et al., 2021c; Levey et al., 2022; Lewis et al., 2023; Li et al., 2021; Lin et al., 2022; Vaidya et al., 2023; Zhao et al., 2022).

NeuroMark was developed as an accessible and flexible framework, with the only core requirement being input fMRI data spatially normalized to the standard adult MNI space (Friston, 2003). However, this assumption raises an important methodological question: when data are acquired from populations whose brain morphology differs markedly from that of young adults—such as infants, children, adolescents, or older adults—does direct alignment to an adult template introduce systematic spatial distortion or bias? This concern is highly relevant because neuroimaging studies frequently recruit age-defined cohorts, reflecting the distinct developmental trajectories, disease incidence patterns, and neurobiological mechanisms that characterize different stages of the lifespan. Ensuring that the normalization approach does not compromise downstream feature extraction is therefore critical for valid cross-age comparisons and reliable biomarker discovery.

Given the broad adoption of both EPInorm and T1toEPInorm within contemporary fMRI preprocessing pipelines, the present study systematically compared these two approaches across multiple large-scale cohorts spanning distinct developmental and aging phases. Previous evidence indicates that EPInorm can perform on par with, or in some instances exceed, normalization strategies based on individualized T1-weighted images, yielding enhanced inter-subject registration consistency and greater effective statistical power (Calhoun et al., 2017). Building on these findings, the current work shifts emphasis from individualized anatomy to population-based structural templates within the T1toEPInorm strategy. This design enables assessment of whether registration to cohort-representative T1 templates—rather than direct EPI-to-adult-template normalization—produces significant differences in NeuroMark-derived features across the lifespan.

Our findings demonstrate that EPInorm and T1toEPInorm yield highly comparable NeuroMark-derived features across all examined cohorts, indicating that a population-specific template during spatial normalization is optional for the effective implementation of the NeuroMark framework. These results contribute to an ongoing methodological debate concerning whether fMRI data should be normalized to population-specific templates rather than to a standard adult reference (Hsu et al., 2007; Sanchez et al., 2012; Weng et al., 2015; Wilke et al., 2002b). This debate is motivated by several neurobiological and analytical considerations. Although standard adult templates—typically derived from young, healthy adults—are favored due to their accessibility, well-established anatomical conventions, and seamless integration into widely used neuroimaging toolkits (e.g., SPM, FSL, AFNI), reliance on a single adult template presumes morphological similarity across diverse populations, an assumption often violated in practice. Neuroimaging research has consistently shown pronounced age-related changes in brain morphology, tissue contrast, and functional topography, encompassing rapid neurodevelopment in infancy and childhood, protracted maturation through adolescence, and progressive cortical and subcortical atrophy in aging (Lockhart and DeCarli, 2014; Sussman et al., 2016; Tamnes et al., 2010). Such developmental and aging-related heterogeneity can introduce systematic normalization errors when non-adult brains are co-registered to an adult template, including boundary misregistration, suboptimal alignment of functional networks, and potential biases in downstream feature extraction.

However, population-specific normalization presents significant practical and scientific trade-offs, the most important of which is the reduced comparability across datasets (Carmon et al., 2020; Pappas et al., 2021). Variability in reference spaces and registration procedures can introduce systematic differences between studies, making cross-dataset harmonization more complex and potentially undermining large-scale integrative analyses. This issue is particularly relevant for NeuroMark, which was explicitly designed to generate feature representations that are comparable across subjects, studies, and datasets. Its goal is to support reproducible population-level neuroscience and to investigate both the commonalities and disorder-specific uniqueness of brain abnormalities. The introduction of age- or cohort-specific templates during normalization may unintentionally modify the extracted NeuroMark features, complicating interpretation and limiting generalizability. Moreover, several lines of evidence suggest that modest variability in anatomical alignment does not necessarily lead to meaningful differences in functional analyses, especially when using higher-order, data-driven models like ICA or network-level analyses. Inter-individual anatomical variation and minor discrepancies in spatial registration have a limited impact on functional localization and activation patterns (Buckner et al., 2004; Thirion et al., 2007). Consistent with this body of literature, our findings demonstrate that NeuroMark features—including ICN spatial maps, time course dynamics, and FNC—remain highly stable across different normalization strategies. This might suggest ICA-derived functional metrics are resilient to differences in preprocessing methods, as data-driven higher-order models can effectively accommodate modest anatomical variability while preserving functional interpretability (Calhoun et al., 2009; Noble et al., 2019; Zuo et al., 2019).

Geometric distortion correction constitutes a critical methodological consideration when comparing direct EPI-based normalization with T1-driven normalization pipelines. Unlike EPInorm, T1-based approaches do not inherently compensate for susceptibility-induced distortions or signal dropout artifacts, which predominantly affect EPI data and can lead to suboptimal cross-modal alignment if not adequately corrected before registration. Because these distortions are often pronounced near air–tissue interfaces and vary substantially across individuals, contemporary preprocessing frameworks increasingly incorporate dedicated distortion-mapping acquisitions (e.g., field maps) (In et al., 2015) or interleaved reverse–phase-encoded images (Van Essen et al., 2012) to enhance the accuracy of EPI distortion correction. In the present study, we employed rigorous distortion- and signal–dropout– correction procedures before spatial normalization to minimize potential misalignment between T1 and EPI modalities. Under these optimized conditions, EPInorm and T1toEPInorm produced highly concordant NeuroMark-derived spatial maps, intrinsic network time courses, and functional connectivity estimates. These findings reinforce the robustness of NeuroMark-derived imaging features to differences in normalization strategy, even in cohorts whose brain morphology diverges substantially from the adult reference. We further observed that, across large-scale intrinsic networks, sensory networks exhibited the highest robustness to normalization strategy, whereas subcortical and default mode networks showed comparatively greater sensitivity—although still demonstrating very high cross-method consistency. Based on these results, we recommend EPInorm for studies involving heterogeneous or lifespan-diverse cohorts, where the use of age-specific structural templates could introduce unintended biases. For studies with age-homogeneous samples or analyses focused more narrowly on subcortical or default mode systems, both EPInorm and T1toEPInorm appear suitable and may be used interchangeably for examining the reliability of NeuroMark feature extraction.

Several limitations of the current study warrant consideration and provide directions for future research. First, our investigation focused exclusively on the reliability and robustness of static NeuroMark-derived features. However, NeuroMark can also extract a broad spectrum of dynamic metrics, such as dynamic functional network connectivity (dFNC) (Fu et al., 2025), dynamic amplitude of low-frequency fluctuations (dALFF) (Fu et al., 2021a, 2018), and spatially dynamic network configurations (Iraji et al., 2024). Given the increasing recognition of the brain’s intrinsically dynamic organization (Hutchison et al., 2013), an important next step is to determine whether these dynamic features also exhibit comparable robustness across varying age cohorts and normalization strategies. Such work would further strengthen the generalizability of the NeuroMark framework. Second, accumulating evidence indicates substantial heterogeneity in brain organization across distinct neuropsychiatric and neurological disorders, reflecting disorder-specific pathophysiological mechanisms (Qiu et al., 2011; Raji et al., 2009; Rudie et al., 2013; Strakowski et al., 1999; Zhang et al., 2018). Although NeuroMark has demonstrated strong adaptability and reproducibility across diverse patient populations, it remains unknown whether incorporating disorder-specific templates during spatial normalization could enhance sensitivity to disease-related alterations. Future studies could explore the development of disorder-informed normalization templates for EPI data and examine whether disease-associated morphological deviations influence NeuroMark-based feature extraction during alignment to standard space. Third, recent work has extended the NeuroMark framework beyond fMRI by constructing multimodal templates applicable to structural MRI (sMRI) (Fu et al., 2024), diffusion MRI (dMRI) (Fu et al., 2024), and PET imaging (Eierud et al., 2024). Analogous to fMRI, these modalities require spatial normalization before NeuroMark analysis. Therefore, assessing how different normalization strategies affect multimodal NeuroMark feature extraction will be essential for establishing consistent and modality-generalizable preprocessing pipelines for NeuroMark.

## Ethics Statement

All infant datasets analyzed in this study were collected under research protocols approved by the Emory University Institutional Review Board (IRB). Written informed consent was obtained from the legal guardians of all participating infants at the time of enrollment. The Human Connectome Project (HCP) datasets used in this work consist of open-access, fully de-identified human imaging data; therefore, no additional institutional ethical review was required.

## Data and Code Availability Statement

The HCP Development and HCP Aging datasets analyzed in the present study are available through the National Institute of Mental Health Data Archive (NDA; https://nda.nih.gov/) and can be accessed upon submission of a data-use application and approval from the ABCD consortium. The NeuroMark templates are publicly accessible via the TReNDS Center website (https://trendscenter.org/data/) and the associated GitHub repository (https://github.com/trendscenter/gift/tree/master/GroupICAT/icatb/icatb_templates). The NeuroMark framework has been fully integrated into the Group ICA Toolbox (GIFT; https://trendscenter.org/software/gift/). Additional MATLAB scripts used in the present analyses are available from the corresponding author upon reasonable request.

## Additional Information

Supplementary Materials are available.

