## Supplementary Material for "Robustness of NeuroMark-Derived Functional Networks to fMRI Spatial Normalization Across the Human Lifespan"

### List of Contents

### Emory Infant Cohorts

The infant cohorts were drawn from ongoing longitudinal studies at the Marcus Autism Center (Atlanta, GA, USA) investigating neurodevelopment in infants at high versus low familial likelihood for autism spectrum disorder (ASD). Infants were enrolled into one of two groups: (1) a high-likelihood group, defined by having a first-degree relative with ASD, and (2) a low-likelihood comparison group, defined by no family history of ASD within three degrees. Exclusion criteria for both groups included gestational age <34 weeks, significant pre-/perinatal complications, major sensory impairments, non-febrile seizures, known genetic syndromes, or MRI contraindications. The Emory University Institutional Review Board approved all procedures.

Each infant was scheduled for up to three pseudorandomly timed MRI sessions between birth and 9 months of age. Imaging was performed during natural sleep on a 3T Siemens Tim Trio or 3T Siemens Prisma system (32-channel head coil) at the Emory Center for Systems Imaging. Infants were swaddled and soothed to sleep before scanning and were continuously monitored via an MRI-compatible camera and in-ear optical microphones. Scanner noise was kept below 80 dBA using pediatric sound-attenuating headphones and a custom acoustic hood, with gradually increasing white noise delivered before the first sequence to mask sequence onset. A trained researcher remained in the scanner room, and scanning was stopped upon any signs of awakening or elevated sound levels.

Resting-state functional MRI data were acquired using multiband BOLD echo-planar imaging. On the Tim Trio, acquisition parameters were: TR = 720 ms, TE = 33 ms, flip angle = 53°, FOV = 208 × 208 mm, 72 slices, and 570 volumes (6 min 50.4 s). On the Prisma, parameters were: TR = 800 ms, TE = 37 ms, flip angle = 52°, FOV = 208 × 208 mm, 72 slices, and 420 volumes (5 min 36 s). For distortion correction, scans were collected in both anterior-to-posterior and posterior-to-anterior phase-encoding directions, yielding two rs-fMRI runs per session.

### HCP Development Cohorts

Childhood and adolescence constitute periods of profound neurobiological maturation, during which the human brain undergoes extensive structural and functional reorganization (Fair et al., 2009; Giedd et al., 1999). These neural changes co-occur with marked physical, cognitive, emotional, and social development, yet the mechanisms by which evolving brain architecture shapes emerging behavior—and how atypical developmental trajectories relate to disrupted neurocircuitry—remain incompletely understood (Fox et al., 2010). The Human Connectome Project in Development (HCP-D) addresses this gap by characterizing normative brain development and its variability in childhood and adolescence. The HCP-D cohort includes cross-sectional structural and functional MRI data from 652 healthy participants, complemented by extensive demographic and behavioral assessments.

HCP-D data were acquired using the Lifespan HCP imaging protocol, which closely parallels the Young Adult HCP protocol with minor modifications to accommodate scanner hardware differences, reduce total scanning time, and address practical challenges inherent to scanning younger and older populations (Harms et al., 2018). All data were collected on Siemens 3T Prisma systems equipped with 80 mT/m gradients (200 T/m/s slew rate) and a 32-channel head coil to support high multiband acceleration. Resting-state fMRI was acquired using a 2D multiband gradient-echo EPI sequence (MB = 8; TR/TE = 800/37 ms; flip angle = 52°) with 2.0-mm isotropic voxels across 72 oblique-axial slices. Each session included two functional runs with opposite phase-encoding directions (anterior–posterior and posterior–anterior) to mitigate bias from susceptibility-induced distortions. Each run lasted 6 minutes 41 seconds (488 frames), yielding high-quality functional connectivity data suitable for lifespan analyses.

### HCP Aging Cohorts

Complementing the developmental arm of the Lifespan HCP initiative, the Human Connectome Project in Aging (HCP-A) extends this framework to investigate normative variation in brain structure and function from midlife through older adulthood (Bookheimer et al., 2019). Converging evidence indicates that large-scale functional brain organization changes systematically across the adult lifespan, and characterizing these age-related alterations is essential for distinguishing typical aging from the neural perturbations associated with late-life neurodegenerative and psychiatric disorders (Erdő et al., 2017; Mrak et al., 1997). The HCP-A study was therefore designed to map healthy aging trajectories of brain circuitry and to provide a reference for understanding individual differences that may confer vulnerability to conditions such as dementia.

Using an experimental design parallel to HCP-D, the HCP-A dataset comprises cross-sectional resting-state fMRI, structural imaging, and extensive demographic and behavioral assessments from 725 healthy adults aged 36–100 years. Data acquisition followed the Lifespan HCP imaging protocol, which maintains substantial alignment with the Young Adult HCP protocol while incorporating minor adjustments necessitated by hardware differences, reduced per-participant scan burden, and the practical constraints of scanning both younger and older populations. All scans were collected on Siemens 3T Prisma systems equipped with 80 mT/m gradients (200 T/m/s slew rate) and a 32-channel head coil enabling high multiband acceleration. Resting-state fMRI was acquired using a 2D multiband gradient-echo EPI sequence (MB = 8; TR/TE = 800/37 ms; flip angle = 52°) with 2.0-mm isotropic voxels across 72 oblique-axial slices. As in HCP-D, functional imaging consisted of paired runs with opposing phase-encoding directions (anterior–posterior and posterior–anterior) to mitigate susceptibility-induced distortions. Each run lasted 6 minutes and 41 seconds (488 frames), producing high-quality, harmonized functional connectivity data suitable for lifespan analyses.

### Preprocessing of fMRI

Preprocessing of the resting-state fMRI datasets followed a rigorously standardized workflow. To ensure signal stabilization and reduce transient magnetization effects, the first ten volumes of each run were discarded. Head motion, an especially prominent concern in infant neuroimaging, was corrected using FSL’s *mcflirt*, with all volumes rigidly realigned to the first frame to provide a consistent reference for downstream processing. Susceptibility-induced field inhomogeneities, which can produce substantial geometric distortions in EPI-based acquisitions, were estimated from the single-band calibration scans collected with opposing anterior–posterior and posterior–anterior phase-encoding directions. These dual-polarity field maps were used to compute the off-resonance field, which was subsequently applied to correct distortions in the multiband BOLD data. Slice-timing correction was performed after distortion compensation to account for temporal offsets introduced by the multiband acquisition scheme. Spatial normalization was carried out using either the EPInorm or T1toEPInorm approach. Both strategies register the infant data to the standardized adult EPI template, thereby ensuring compatibility with the NeuroMark framework. EPInorm applies direct EPI-to-EPI registration, whereas T1toEPInorm first leverages an age-specific T1 anatomical template before transforming into adult space, providing an alternative pathway for harmonizing infant scans. Finally, fMRI data were resampled to 3 × 3 ×3 mm3 isotropic voxels using SPM, and the normalized images were smoothed with a 6-mm full-width at half-maximum Gaussian kernel to enhance signal-to-noise ratios, accommodate inter-individual anatomical variability, and support robust estimation of large-scale functional network structure in subsequent analyses.

### Normalization to Standard MNI Space

In this study, we evaluated two spatial normalization strategies for aligning different EPI data to the standard MNI space (Penny et al., 2011), a prerequisite for applying the NeuroMark framework in a manner that ensures cross-subject and cross-cohort comparability. The first approach, EPInorm, is a single-step procedure in which each subject’s EPI image is directly registered to the SPM young-adult EPI template. This method maintains the data within a consistent contrast domain throughout the registration process, thereby avoiding potential mismatches that arise when aligning EPI data to structural images with markedly different tissue contrast. EPInorm has served as the primary normalization strategy in the majority of our prior work. Accumulating evidence shows that direct EPI-to-template registration performs as well as, and in several cases better than, T1-based normalization pipelines—even when both approaches incorporate distortion correction (Calhoun et al., 2017). Advantages reported in the literature include reduced across-subject variability, improved alignment of functional boundaries, and enhanced statistical sensitivity in voxelwise analyses. These properties make EPInorm a practical and robust choice for large-scale functional connectivity studies, particularly when consistent alignment of intrinsic functional networks is a central aim.

The second normalization strategy was a two-stage, anatomically informed procedure that leveraged age-specific high-resolution structural templates to improve the accuracy of spatial normalization into the common MNI space. In this T1toEPInorm approach, functional EPI data were first aligned to an age-appropriate T1-weighted anatomical template, and the resulting intermediate representation was then transformed into the adult MNI EPI space. This hierarchical normalization framework exploits age-specific anatomical priors to reduce registration bias and enhance cross-subject alignment, consistent with widely adopted practices in large-scale developmental and lifespan neuroimaging pipelines. For the infant cohort, considering the rapid brain maturation during infancy, EPI images were first normalized to a densely sampled series of age-specific T1 templates spanning birth through nine months, derived from the UNC/UMN Baby Connectome Project (BCP) (Chen et al., 2022). Each infant’s scan was matched to the T1 template corresponding to their age at acquisition. Using SPM’s standard normalization algorithm, the native-space fMRI data were warped into the selected BCP T1 template. Subsequently, a second transformation was applied to map the age-normalized data into the SPM adult EPI MNI template, thereby ensuring compatibility with the NeuroMark spatial priors. For the developmental cohort, we employed unbiased pediatric T1 atlases generated by the NeuroImaging & Surgical Technologies Laboratory at McGill University (Fonov et al., 2011, 2009). These atlases were designed to capture normative age-related variation in neuroanatomy across childhood and adolescence. A single representative pediatric T1 template (ages 4.5–18.5 years) was selected for coregistration, and the same two-step normalization pipeline was then applied. For the aging cohort, spatial alignment was anchored using the Multichannel Illinois Institute of Technology and Rush University Aging (MIITRA) atlas (Wu et al., 2023), a high-quality T1 template constructed using advanced, multimodal template-building techniques from healthy older adults. Similarly, a single T1 template served as the anatomical reference for the first normalization stage. Across all cohorts, EPI data normalized by the EPInorm and the T1toEPInorm were subsequently entered into the NeuroMark pipeline to enable systematic comparison of normalization strategies and their downstream effects on functional network estimation.

### Mask Generating for NeouroMark Framework

For each cohort, cohort-specific brain masks were generated using a spatial–similarity–driven procedure designed to ensure high consistency across individuals and to provide robust constraints for subsequent NeuroMark analyses. Mask generation proceeded in several stages. First, an initial brain mask was estimated for each scan using the first volume of the fMRI time series. Voxels were classified as brain tissue if their intensity exceeded 90% of the mean signal intensity across all voxels within that volume; all other voxels were set to zero. This step yielded a binary mask for every scan. Second, a provisional group-level mask was constructed by retaining only those voxels that were included in at least 90% of the individual masks. To evaluate the reliability of each mask relative to this group reference, we computed spatial correlations along three dimensions: (i) whole-brain correlation across all voxels within the preliminary group mask, (ii) top-slice correlation using only the upper ten axial slices, and (iii) bottom-slice correlation using the lower ten axial slices. These regional correlations were designed to capture potential inconsistencies caused by partial-brain coverage or low signal fidelity at the superior or inferior boundaries of the acquisition. A scan was deemed suitable for mask construction only if it exceeded prespecified thresholds on all three metrics (whole-brain r > 0.8, top r > 0.75, bottom r > 0.55). This quality-control step ensured that only high-fidelity, spatially reliable masks contributed to the final group representation. Finally, a refined group mask was recomputed using only the subset of scans meeting these criteria, yielding a high-quality cohort-specific mask optimized for guiding NeuroMark component estimation.

Because the masks derived from EPInorm- and T1toEPInorm-processed data exhibited minimal differences, and to maintain fairness in downstream comparisons between normalization strategies, we adopted the group mask derived from the T1toEPInorm pipeline for all subsequent analyses.

### NeuroMark Framework and Feature Extraction

We applied a hybrid, data-driven analytic framework—NeuroMark(Du et al., 2020; Fu et al., 2024)—to the resting-state fMRI datasets to derive subject-specific network features for downstream cross-strategies comparison analyses. NeuroMark integrates the advantages of traditional atlas-based parcellation with the flexibility of ICA by combining a robust, population-derived spatial network template with spatially constrained ICA decomposition. This template-guided strategy mitigates well-recognized limitations of purely atlas-dependent methods—such as arbitrary boundary definitions and restricted adaptability to individual functional organization—as well as those of unconstrained ICA, which can yield components that lack anatomical correspondence across participants and studies. By imposing empirically validated spatial priors, NeuroMark facilitates the extraction of ICNs that are both biologically interpretable and directly comparable across individuals, cohorts, and acquisition sites, thereby enhancing reproducibility, generalizability, and sensitivity to inter-individual variability.

In the present study, we utilized the NeuroMark 1.0 functional template as the reference space. This template comprises 53 highly reproducible ICNs, each exhibiting peak activation within canonical gray-matter territories and characterized by dominant low-frequency fluctuations in their time courses. The ICNs are organized into seven functional domains—sensorimotor, visual, cognitive control, default mode, auditory, subcortical, and cerebellar—following established neuroanatomical and functional taxonomy. The spatial maps of these components are presented in Fig. S1, and full network labels and domain assignments are provided in Table S1.

For each dataset, ICN spatial maps and their associated time courses were estimated using group-information-guided ICA (GIG-ICA) (Du and Fan, 2013), a back-reconstruction algorithm that jointly optimizes (i) the statistical independence of single-subject components and (ii) their spatial correspondence to the NeuroMark template. Relative to conventional ICA back-reconstruction procedures, GIG-ICA has demonstrated superior reconstruction fidelity and higher intra-class correlation coefficients for single-subject ICNs, reflecting improved stability and reliability (Du et al., 2016). Furthermore, GIG-ICA enhances sensitivity to inter-individual and clinical variation in network architecture, making it particularly well-suited for identifying functional biomarkers in both normative and clinical populations.

The NeuroMark framework, together with the 1.0 functional template, has been extensively validated and widely deployed across large-scale neuroimaging studies. It has consistently demonstrated robust performance in detecting reproducible intrinsic brain networks and in characterizing functional dysconnectivity across diverse neuropsychiatric and neurodevelopmental conditions.

### NeuroMark Functional 1.0 Template

The NeuroMark functional 1.0 template was derived from two large, independent resting-state fMRI cohorts comprising a combined sample of 1,828 healthy adults: the Human Connectome Project (HCP) (Van Essen et al., 2012) and the Genomics Superstruct Project (GSP) (Buckner et al., 2014). For each dataset, a high–model-order group independent component analysis (ICA; model order = 100) was performed to decompose the data into spatially independent components (ICs). Cross-dataset correspondence was then established by computing voxel-wise spatial correlations between ICs from the HCP and GSP decompositions. Component pairs exhibiting a spatial correlation ≥ 0.40 were retained as reproducible features. Although correlations ≥ 0.25 have been shown to reflect statistically significant cross-dataset correspondence (p < 0.005, corrected) (Smith et al., 2009), the more stringent threshold was adopted to ensure that only highly robust and reliably replicating ICs were selected. Matched IC pairs were subsequently classified as putative ICNs or noise components based on established criteria, including the anatomical plausibility and gray-matter localization of peak activations, as well as the predominance of low-frequency fluctuations in their associated time courses. Among the IC pairs identified as bona fide ICNs, the GSP-derived components, which exhibited comparatively reduced noise and more stable spatial profiles, were selected as the networks (**Table SI, Figure S1**) for building the NeuroMark functional 1.0 template (Du et al., 2020).

Table SI. Peak Coordinates of Networks from NeuroMark Functional 1.0 Template

| **Intrinsic Connectivity Networks** | **X** | **Y** | **Z** |
| --- | --- | --- | --- |
| **Subcortical Domain (SC)** | | | |
| Caudate (69) | 6.5 | 10.5 | 5.5 |
| Subthalamus/hypothalamus (53) | -2.5 | -13.5 | -1.5 |
| Putamen (98) | -26.5 | 1.5 | -0.5 |
| Caudate (99) | 21.5 | 10.5 | -3.5 |
| Thalamus (45) | -12.5 | -18.5 | 11.5 |
| **Auditory Domain (AUD)** | | | |
| Superior temporal gyrus ([STG], 21) | 62.5 | -22.5 | 7.5 |
| Middle temporal gyrus ([MTG], 56) | -42.5 | -6.5 | 10.5 |
| **Sensorimotor Domain (SM)** | | | |
| Postcentral gyrus ([PoCG], 3) | 56.5 | -4.5 | 28.5 |
| Left postcentral gyrus ([L PoCG], 9) | -38.5 | -22.5 | 56.5 |
| Paracentral lobule ([ParaCL], 2) | 0.5 | -22.5 | 65.5 |
| Right postcentral gyrus ([R PoCG], 11) | 38.5 | -19.5 | 55.5 |
| Superior parietal lobule ([SPL], 27) | -18.5 | -43.5 | 65.5 |
| Paracentral lobule ([ParaCL], 54) | -18.5 | -9.5 | 56.5 |
| Precentral gyrus ([PreCG], 66) | -42.5 | -7.5 | 46.5 |
| Superior parietal lobule ([SPL], 80) | 20.5 | -63.5 | 58.5 |
| Postcentral gyrus ([PoCG], 72) | -47.5 | -27.5 | 43.5 |
| **Visual Domain (VS)** | | | |
| Calcarine gyrus ([CalcarineG], 16) | -12.5 | -66.5 | 8.5 |
| Middle occipital gyrus ([MOG], 5) | -23.5 | -93.5 | -0.5 |
| Middle temporal gyrus ([MTG], 62) | 48.5 | -60.5 | 10.5 |
| Cuneus (15) | 15.5 | -91.5 | 22.5 |
| Right middle occipital gyrus ([R MOG], 12) | 38.5 | -73.5 | 6.5 |
| Fusiform gyrus (93) | 29.5 | -42.5 | -12.5 |
| Inferior occipital gyrus ([IOG], 20) | -36.5 | -76.5 | -4.5 |
| Lingual gyrus ([LingualG], 8) | -8.5 | -81.5 | -4.5 |
| Middle temporal gyrus ([MTG], 77) | -44.5 | -57.5 | -7.5 |
| **Cognitive-control Domain (CC)** | | | |
| Inferior parietal lobule ([IPL], 68) | 45.5 | -61.5 | 43.5 |
| Insula (33) | -30.5 | 22.5 | -3.5 |
| Superior medial frontal gyrus ([SMFG], 43) | -0.5 | 50.5 | 29.5 |
| Inferior frontal gyrus ([IFG], 70) | -48.5 | 34.5 | -0.5 |
| Right inferior frontal gyrus ([R IFG], 61) | 53.5 | 22.5 | 13.5 |
| Middle frontal gyrus ([MiFG], 55) | -41.5 | 19.5 | 26.5 |
| Inferior parietal lobule ([IPL], 63) | -53.5 | -49.5 | 43.5 |
| Left inferior parietal lobue ([R IPL], 79) | 44.5 | -34.5 | 46.5 |
| Supplementary motor area ([SMA], 84) | -6.5 | 13.5 | 64.5 |
| Superior frontal gyrus ([SFG], 96) | -24.5 | 26.5 | 49.5 |
| Middle frontal gyrus ([MiFG], 88) | 30.5 | 41.5 | 28.5 |
| Hippocampus ([HiPP], 48) | 23.5 | -9.5 | -16.5 |
| Left inferior parietal lobule ([L IPL], 81) | -47.5 | 5.5 | 22.5 |
| Middle cingulate cortex ([MCC], 37) | -15.5 | 20.5 | 37.5 |
| Inferior frontal gyrus ([IFG], 67) | 39.5 | 44.5 | -0.5 |
| Middle frontal gyrus ([MiFG], 38) | -26.5 | 47.5 | 5.5 |
| Hippocampus ([HiPP], 83) | -24.5 | -36.5 | 1.5 |
| **Default-mode Domain (DM)** | | | |
| Precuneus (32) | -8.5 | -66.5 | 35.5 |
| Precuneus (40) | -12.5 | -54.5 | 14.5 |
| Anterior cingulate cortex ([ACC], 23) | -2.5 | 35.5 | 2.5 |
| Posterior cingulate cortex ([PCC], 71) | -5.5 | -28.5 | 26.5 |
| Anterior cingulate cortex ([ACC], 17) | -9.5 | 46.5 | -10.5 |
| Precuneus (51) | -0.5 | -48.5 | 49.5 |
| Posterior cingulate cortex ([PCC], 94) | -2.5 | 54.5 | 31.5 |
| **Cerebellar Domain (CB)** | | | |
| Cerebellum ([CB], 13) | -30.5 | -54.5 | -42.5 |
| Cerebellum ([CB], 18) | -32.5 | -79.5 | -37.5 |
| Cerebellum ([CB], 4) | 20.5 | -48.5 | -40.5 |
| Cerebellum ([CB], 7) | 30.5 | -63.5 | -40.5 |

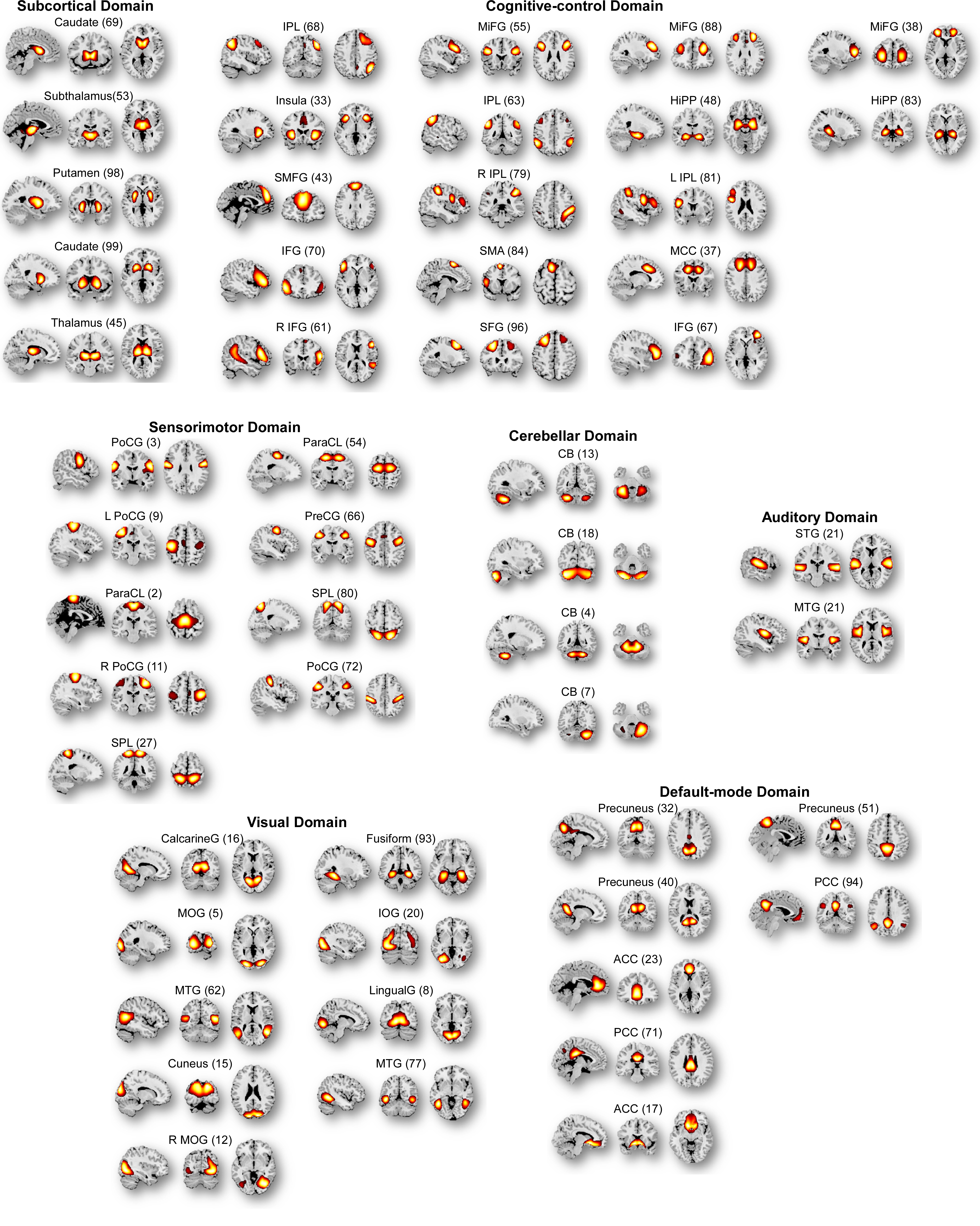

Figure S1. **Spatial topographies of intrinsic connectivity networks (ICNs) from the NeuroMark 1.0 functional template.** The 53 ICNs are organized into seven canonical functional domains based on established anatomical and functional ontologies. Spatial maps were generated by computing one-sample *t*-statistics across single-scan ICN maps and visualized using a threshold of |*t*| > 10. For each network, sagittal, coronal, and axial views are displayed at the voxel exhibiting the maximal *t*-value within clusters exceeding 3 cm³ in volume, highlighting the core topography of each ICN.
